# Inclusion of Hsp70 co-chaperone and RNA binding activities facilitate optimal function of spliceosomal disassembly factor Cwf23 in intron-rich *Schizosaccharomyces pombe*

**DOI:** 10.1101/2024.06.27.600868

**Authors:** Kirpa Yadav, Sandeep Raut, Meghna G Roy, S Gopika, Minita Desai, R. Balashankar, Shravan Kumar Mishra, Chandan Sahi

## Abstract

Cwc23 is an essential J-domain protein that collaborates with the disassembly factor Ntr1 to facilitate spliceosomal disassembly. Although Cwc23 orthologs have been identified in spliceosomal extracts of many eukaryotes, their functionality in intron-rich eukaryotes remains largely unexplored. Here, we investigate the functionality of Cwf23, an ortholog of Cwc23 in *Schizosaccharomyces pombe*. Our study reveals that while the interaction between Cwf23 and *Sp*Ntr1 is conserved, it is not essential in *S. pombe*. Additionally, the RNA recognition motif (RRM) in Cwf23 is crucial for its function, as mutations in the RRM affect both growth and pre-mRNA splicing. Unlike its budding yeast counterpart, the J-domain of Cwf23 is indispensable, as J-domain mutants cannot support cell viability. These findings suggest a new paradigm in which the presence of a functional RRM and the essential nature of the J-domain underscore the increased requirements for an RNA-binding Hsp70 co-chaperone machinery in spliceosomal remodelling in “intron-rich” eukaryotes.

## Introduction

Spliceosome is held together by numerous protein-protein, RNA-protein, interactions. Throughout the process of splicing, the spliceosome undergoes a large number of conformational and compositional changes brought about by the formation and disruption of RNA–protein and protein-protein interactions (Fabrizio et al., 2009; Hoskins & Moore, 2012; Staley & Guthrie, 1998; Will & Lührmann, 2011). Each round of splicing includes assembly, activation, catalysis and disassembly of the spliceosomal components which are then recycled for a fresh round of splicing (Hoskins & Moore, 2012).

Spliceosomal disassembly causes the post-catalytic deconstruction of the spliceosome to release mature mRNA, snRNPs, and splicing factors and recycle the latter two for successive rounds of pre-mRNA splicing. Best characterized in budding yeast, this process is mediated by many spliceosomal proteins, including the DEAH-box RNA helicases Prp22 and Prp43. Prp22 translocates along the mature mRNA in a 3’-to-5’ direction and supports its release from the intron-lariat spliceosome (ILS) in the initial step of this disassembly (Company et al., 1991; Schwer, 2008). Prp43 then facilitates the separation of the IL (Intron Lariat) RNA from the ILS (Arenas & Abelson, 1997; Martin et al., 2002; Tsai et al., 2005). Simultaneously, the remaining spliceosomal core containing the U2/U6 and U5 snRNPs is also released (Tsai et al., 2007). Prp43 interacts with two other splicing factors Ntr1 (also known as Spp382) and Ntr2 forming the <u>n</u>ineteen-<u>r</u>elated (NTR) complex (Boon et al., 2006; Tsai et al., 2005). Ntr1 acts as a scaffold protein in this complex which interacts with Prp43 through its N-terminal G-patch domain and with Ntr2 through its C-terminal domain (Aravind & Koonin, 1999; Tsai et al., 2005). Ntr1 is required to stimulate the otherwise weak helicase activity of Prp43 to bring about the spliceosome disassembly (Tanaka et al., 2007; Tsai et al., 2007). Ntr1 and Ntr2 act as cofactors for Prp43, facilitating the removal of spliceosome intermediates at different stages of the spliceosomal cycle before disassembly (Chen et al., 2013; Mayas et al., 2010; Pandit et al., 2006).

The C-terminal of Ntr1 also interacts with a J-domain protein (JDP), Cwc23, whose precise role in spliceosome remodelling remains largely elusive. Cwc23 is an essential JDP conserved across eukaryotes (Raut et al., 2019). JDPs are co-chaperones of heat shock protein 70 (Hsp70s). All JDPs have a signature J-domain which is critical for stimulating the ATPase activity of Hsp70s. Additionally, JDPs also provide functional specificity to their partner Hsp70s (Craig et al., 2006; Kampinga & Craig, 2010). Interestingly, the J-domain, thereby the Hsp70 co-chaperone activity of Cwc23, is completely dispensable for its essential functions. Moreover, a complete deletion of the J-domain does not cause any defects in splicing. Instead, a C-term truncation mutant of Cwc23 resulted in a slow growth phenotype as well as global splicing defects, more specifically, accumulation of intron lariat (IL), a hallmark of malfunctions in the spliceosomal disassembly process (Sahi et al., 2010).

While the conservation of intra-species interaction between Cwc23 and Ntr1 orthologs in more complex eukaryotes, suggests its role in spliceosomal remodeling through the NTR complex, Cwc23 orthologs in intron-rich eukaryotes additionally have an RRM at their C-terminal (Raut et al., 2019). Cwf23, the Cwc23 ortholog in *S. pombe* has a C-terminal RRM capable of binding RNA *in vitro* (Raut et al., 2019). Although the *in vivo* roles of Cwf23 orthologs and their contribution to the process of splicing are still unknown, mutations in DnaJC17, the human ortholog of Cwc23, have been linked to various human disorders including retinal dystrophy, thrombocythemia, congenital hypothyroidism with thyroid dysgenesis and Autism Spectrum Disorder (ASD) (AL Assaf et al., 2014; Hou et al., 2012; Patel et al., 2016).

The present study aimed at deciphering the significance of the RRM and the J-domain of Cwf23 (in *S. pombe*) which is completely dispensable in Cwc23 (*S. cerevisiae*). We show that the RRM, although important for Cwf23 function, is dispensable for interaction with *Sp*Ntr1 and cell viability. Moreover, the interaction between Cwf23 and *Sp*Ntr1 was not crucial as the C-terminal truncations of Cwf23 that did not interact with *Sp*Ntr1 were viable, displaying no observable growth defects. Finally, we find that the J-domain of Cwf23 is essential in *S. pombe*, suggesting an increased requirement of Hsp70-dependent functions of Cwf23 in *S. pombe*. With nearly 45% of genes having introns and more genes with multiple introns, *S. pombe*’s splicing machinery is more complex than *S. cerevisiae* (Neuvéglise et al., 2011). Our results for the first time infer a direct involvement of a specialized Hsp70-JDP system in mediating spliceosome dynamics in complex, intron-rich eukaryotes.

## Materials and methods

### Plasmids construction

The open reading frame (ORF) corresponding to full-length, N-term or C-term deletion of Cwf23 (Fig 1) was PCR amplified using appropriate primers and cloned in pREP3X vectors as BamHI/XhoI fragments for expression in *S. pombe*. Point mutants and internal truncations were created by NEB Q5 Site-Directed Mutagenesis Kit using appropriate primers. Primers were designed using the NEBase Changer tool available online (http://nebasechanger.neb.com/). HA-tagged constructs were generated by adding a sequence corresponding to 1X-HA tag upstream to the ORF of Cwf23 and its mutants. For Y2H analysis, ORF for Cwf23 was PCR amplified from respective mutant plasmids with primers having attachment B (attB) sites flanking gene-specific nucleotides and cloned first into an entry vector pDONR207 and then into destination yeast two-hybrid vectors pGADT7 or pGBKT7 (Clontech, Takara Bio USA) using Gateway cloning kit (Invitrogen, Thermo Fisher Scientific) as per the manufacturer’s instructions. For bacterial expression, the sequence corresponding to the *ΔRRM* was PCR amplified from the pREP3X_*Cwf23*_*ΔRRM* plasmid and cloned into pGEX6P1 in BamHI and SalI sites. For testing the functionality of the Cwf23 J-domain in budding yeast, ORF corresponding to amino acids 1-165 was PCR amplified from wild-type and *H37Q* mutants and cloned in pRS414 plasmid (Mumberg et al., 1995) under *TEF1* promoter. All constructs were confirmed by restriction digestion followed by sequencing and are listed in Supplementary Table 1.

**Figure 1.**
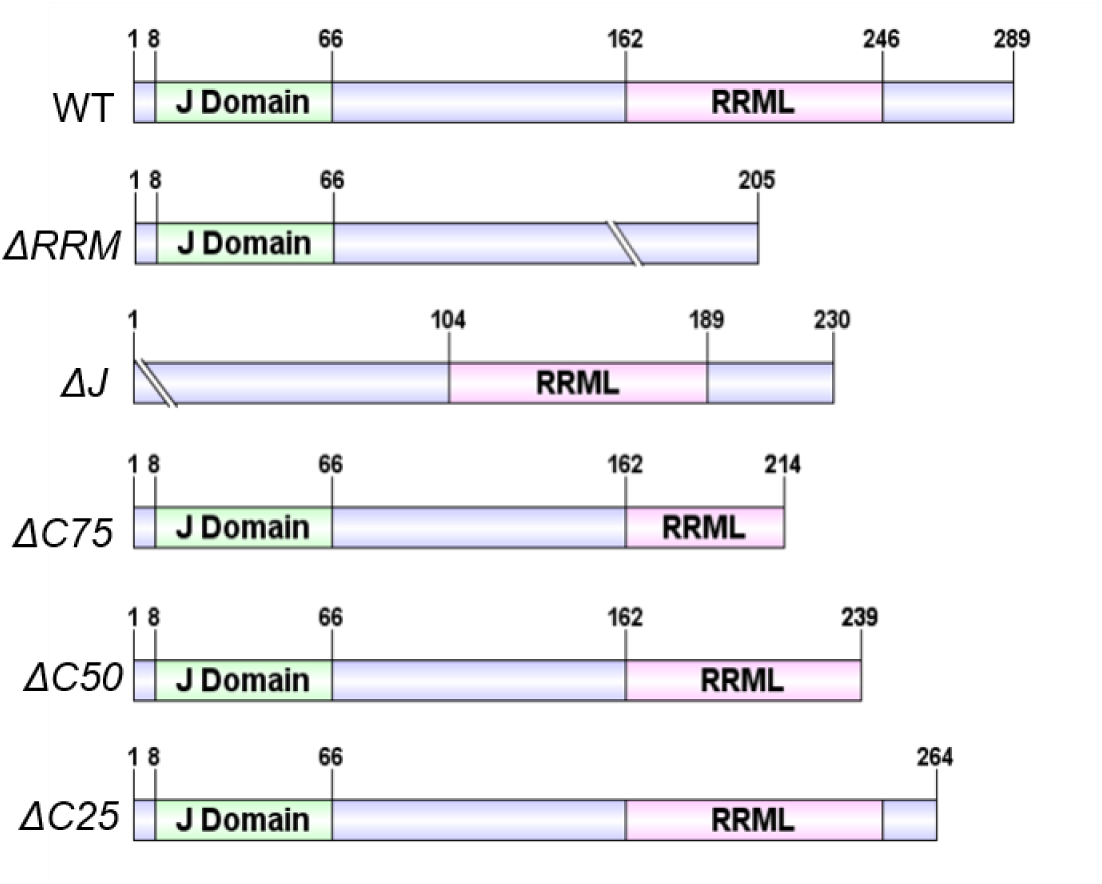
Maps of Cwf23 WT and truncation mutants, showing the regions of deleted residues in the RRM, J-domain and C-term.

### Yeast methods

*Schizosaccharomyces pombe* strain, *cwf23Δ* was generated by first transforming the wild-type *S. pombe* cells (972h(+)) with a *URA3-*based CEN-CWF23 plasmid (Table 1) followed by transformation with the deletion cassette having nourseothricin-resistance gene (*natR*) as the selection marker. The transformants were selected on nat plates. The strain thus obtained was *cwf23Δ* with *URA3* CEN-Cwf23 plasmid. The well-established 5-FOA counter-selection strategy was used for the functional assessment of Cwf23 mutants, in the *cwf23Δ* strain harbouring endogenous copy of Cwf23 on *URA3-*based plasmid. *ydj1Δ S. cerevisiae* strain, used for assessing the functionality of Cwf23 J-domain was described previously (Sahi & Craig, 2007). Yeast transformations were performed using the standard method (Schiestl & Gietz, 1989). Yeast strain *AH109* (Clontech, Takara Bio USA) was used for yeast two-hybrid interaction studies as a tester strain. SD/-Leu/-Trp/-His drop-out has been used as a selective media in this study.

### RNA isolation, cDNA synthesis and splicing assay

Total RNA was isolated from five OD_600_ equivalent cells from a logarithmically growing culture (OD_600_ ∼0.5–0.6) using the acid phenol method, followed by DNase I (NEB, USA) treatment for 30 min at 37°C. 2 μg of total RNA was used to make cDNA using iScript reverse transcriptase (BioRad) as per manufacturer’s instructions. Semi-quantitative PCR was then performed to detect splicing defects. PCR products were resolved and visualised on a 2% agarose gel.

### Protein extraction and western blot analysis

Total protein lysates were prepared as described previously (Knop et al., 1999). Briefly, cells were grown to log phase and harvested by centrifugation at 5000 rcf for 5 minutes. Pellet was resuspended in 1 ml of chilled sterile distilled water. To this, 1.85 N NaOH and 7.5% β-mercaptoethanol was added and incubated on ice for 10 minutes after vortexing. 200 µl of 55% TCA solution was added to it and kept on ice for another 15 minutes. The mixture was centrifuged at 14000 rcf at 4°C for 15 minutes. Supernatant was discarded and centrifuged again to remove the remaining TCA. 50 µl/ OD_600_ HU buffer (8 M urea, 5% SDS, 200 mM Tris pH 6.8, 1 mM EDTA, with bromophenol blue as colouring and pH indicator) with 1.5% DTT was added to the pellet and heated at 65°C for 10 minutes with constant shaking at 1100 rcf. Centrifugation was done at 14000 rcf at RT for 5 minutes and 10 µl of sample was resolved on SDS-PAGE. Following which, proteins were electro-blotted, transferred onto Nitrocellulose membrane and processed for western analysis using appropriate antibodies and chemiluminescence.

### RNA Binding Assays

N-terminal GST Tagged full-length Cwf23 and *ΔRRM* proteins were expressed in *E. coli* Codon Plus cells and affinity purified using Glutathione Agarose Resin (4B-GLU-100, ABT), as described previously (Raut et al., 2019). ∼1000 ng of purified Cwf23, *ΔRRM* and GST proteins were resolved on 12% SDS-PAGE and electro-blotted onto a nitrocellulose membrane. 5 μg of plasmid DNA of *ACT1* was linearized using *SpeI* restriction enzyme (NEB, USA), purified by Phenol:Chloroform extraction and re-suspended in 10 μl of RNase free sterile water (Raut et al., 2019). Riboprobe for the full-length *ACT1* gene was made using “HiScribe™ T7 Quick High Yield RNA Synthesis Kit” (NEB, USA) using Fluorescein labelling mix (Sigma-Aldrich). *In-vitro* transcription was performed as per the manufacturer’s instructions.

Following an on-blot protein renaturation, the *in-vitro* synthesized fluorescein-labelled RNA probe was added to the renatured protein in 2 ml of Binding buffer (20 mM HEPES, pH 7.5, 2.5 mM MgCl2, 1 mM DTT, 5% glycerol dissolved in DEPC-treated water) Hybridization and washings were done exactly as described earlier (Raut et al, 2018). The blot was then incubated with a primary antibody against fluorescein (1:200) at RT followed by an Anti-mouse secondary antibody ((IRDye 800) at 1:20000 dilution for 1 hour at RT. Finally, the blots were washed three times for 5 minutes each using TBST. The blots were then left to dry for 1-2 hours before taking the image using Odyssey 51 Licor imaging system. The fluorescence of the secondary antibody was measured at 800 nm.

## Results

### RRM of Cwf23 is important for cell growth and pre-mRNA splicing but dispensable for interaction with *Sp*Ntr1

Cwc23 is essential in *S. cerevisiae*. While orthologs of Cwc23 have been identified in other eukaryotes, their functional significance, more specifically their role in pre-mRNA splicing has not been studied. In the present study, we aimed to decipher the functional specificity and evolution of Cwf23 in fission yeast. Cwf23 is predicted to have an RRM-like domain, instead of an RRM at the C-terminal as it lacks the characteristic RNP motifs that are present in canonical RRMs. However, the domain topology- βαββαβ is conserved (Raut et al., 2019). RRMs are known to impact protein-protein and protein-RNA interactions. To study the significance of the RRM of Cwf23, we first generated a *cwf23Δ* strain. Since Cwf23 is essential, the viability of this strain was maintained by a *URA-*based plasmid expressing Cwf23 from an endogenous *CWF23* promoter. 5-FOA counter-selection method was employed to assess the functionality of different Cwf23 mutants expressed from a PREP3X plasmid. As expected, *cwf23Δ* strain cells harboring an empty vector could not form colonies on 5-FOA plates, while wild-type Cwf23 could. Our results show that *cwf23Δ* cells expressing the *ΔRRM* mutant could support cell viability but the cells were sick at 30°C as compared to cells expressing wild-type Cwf23 (Fig 2A). N-terminal HA-tagging, followed by western blot analysis, successfully detected both the full-length and truncated Cwf23 proteins, and their expression levels were comparable, suggesting that *Cwf23*_*ΔRRM* was compromised in function (Supplementary Fig 1).

**Figure 2.**
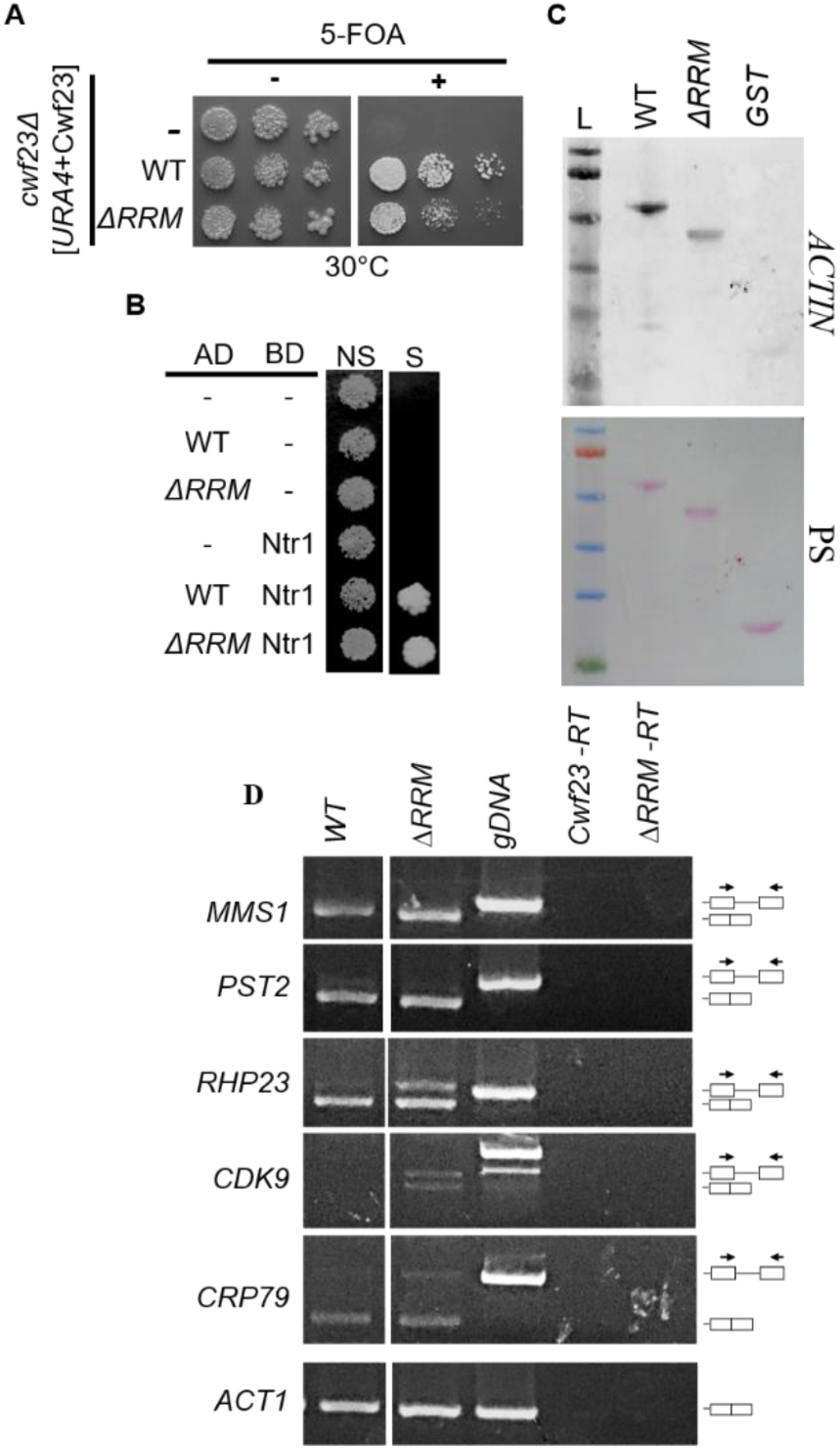
RRM of cwf23 is partially dispensable for cell viability. **A.** Yeast complementation analysis for Cwf23 *ΔRRM*. Equal volumes of ten-fold serial dilutions of *cwf23Δ* cells harboring *URA3 CEN*-Cwf23 plasmid and pREP3x (*LEU*-based plasmid) empty vector (–), Cwf23 (WT) or *ΔRRM* mutant were spotted on media with or without 5-FOA and incubated at 30°C for 3 days. **B.** Deletion of RRM of Cwf23 doesn’t affect interaction with *Sp*Ntr1. Yeast two-hybrid interaction analysis of Cwf23 *ΔRRM* mutant with *Sp*Ntr1. Equal dilutions of two-hybrid tester strain (*AH109*) harbouring *Sp*Ntr1 fused to the *GAL4*-binding domain (BD) and Cwf23 wildtype or *ΔRRM* the fused to the *GAL4* activation domain (AD) or just the empty vectors (-) were spotted on non-selective (NS) or selective (S) medium. Plates were incubated at 30°C for 3 days. Growth on selective medium is indicative of a positive two-hybrid interaction. **C.** In-vitro RNA binding Analysis. Upper panel: ∼1-2 μg of purified Cwf23 (WT), *ΔRRM* mutant & GST were subjected to SDS-PAGE, electro-blotted on to nitrocellulose membrane followed by on-blot renaturation. Blot was then probed with fluorescein-labelled in vitro synthesized transcripts of *S. cerevisiae* ACT1 gene. Lower panel: Ponceau S stained protein bands to show relative amounts of protein electro-blotted. **D.** The RRM of Cwf23 is important for splicing. Agarose gel pictures showing the expression and splicing pattern of selected genes. Total RNA from different strains was used to prepare cDNA. An equal quantity of cDNA was used for PCR with primers specific to each gene flanking the introns. The name of the gene is shown on the left of each panel. Actin primers have been used to verify equal cDNA template for each strain (bottom panel).

Functionality of RNA binding domains (RBDs) depends on their interaction with RNA as well as proteins (Lunde et al., 2007). Previously, we showed that Cwf23 interacts with *Sp*Ntr1 (Raut et al., 2019), so we evaluated whether RRM is required for the interaction between Cwf23 and *Sp*Ntr1. Yeast two-hybrid analysis showed that a complete deletion of RRM did not affect its interaction with *Sp*Ntr1 (Fig 2B), suggesting a role of other regions/domains of Cwf23 in facilitating this interaction. Next, we went ahead to ascertain that RRM is required for RNA binding. For this, an on-blot RNA binding assay was performed (Raut et al., 2019). As shown in Fig 2C, purified *ΔRRM* protein fragment was significantly compromised in binding to *in vitro* synthesized, labelled *ACTIN* RNA as compared to full-length wild-type Cwf23 protein.

Cwf23 has been implicated in pre-mRNA splicing, however, any direct evidence for its function on the spliceosome or splicing is still lacking. Since the *ΔRRM* mutant showed growth defects, we asked if it was also required for pre-mRNA in fission yeast. For this semi-quantitative reverse transcription PCR (RT-PCR) was performed for selected genes. Total RNA was isolated from cells expressing wild-type and *ΔRRM* of Cwf23 and cDNA was synthesized. Multiple genes having one or more introns were selected for this study. Primers were designed for the exonic regions to visualize the spliced (intronless smaller) and unspliced (intron-containing larger) bands in the PCR. Splicing defects were observed in *RHP23* (proteasome-associated ubiquitin receptor) and *CRP79* (poly(A) binding protein) in *ΔRRM* mutant as evidenced by the accumulation of the unspliced mRNAs as compared to the wild-type where only the spliced mRNA bands were detected (Fig 2D). Put together, our data implies that the RRM of Cwf23 is required for optimum growth and pre-mRNA splicing, and the observed defects in *ΔRRM* mutant may be because of impaired RNA binding, an unknown protein-protein interaction, or both.

### Cwf23 C-terminal region is required for interaction with *Sp*Ntr1 but dispensable for cell viability

The most studied ortholog of Cwf23, the budding yeast Cwc23 protein interacts with Ntr1 through its C-terminal region and this interaction is important *in vivo*, as a C-term truncation mutant of Cwc23, *1-225*, is defective in interaction with Ntr1 and also exhibits global splicing defects (Sahi et al., 2010). Since the RRM of Cwf23 was not required for this interaction, we investigated the importance of its C-terminal region. Three C-terminal deletion mutants of Cwf23, deleting the last 25, 50 or 75 amino acids were generated and analyzed for their ability to interact with *Sp*Ntr1 and functionality *in vivo*. While the *ΔC25* mutant interacted with *Sp*Ntr1, *ΔC75* and *ΔC50* did not, in the yeast two-hybrid assays (Fig 3A). Next, we investigated the ability of the C-terminal truncation mutants to substitute for Cwf23 using the 5-FOA counter-selection method. Interestingly, none of these deletions affected Cwf23 function *in vivo* as all the mutants rescued the lethality of *cwf23Δ* strain like wild-type (Fig 3B). Further, these truncations were indistinguishable from the wild-type at different temperatures tested (Fig 3C). Our results suggest that in *S. pombe*, while the C-terminal region of Cwf23 is required for interaction with *Sp*Ntr1, unlike Cwc23, it is dispensable under physiological conditions.

**Figure 3.**
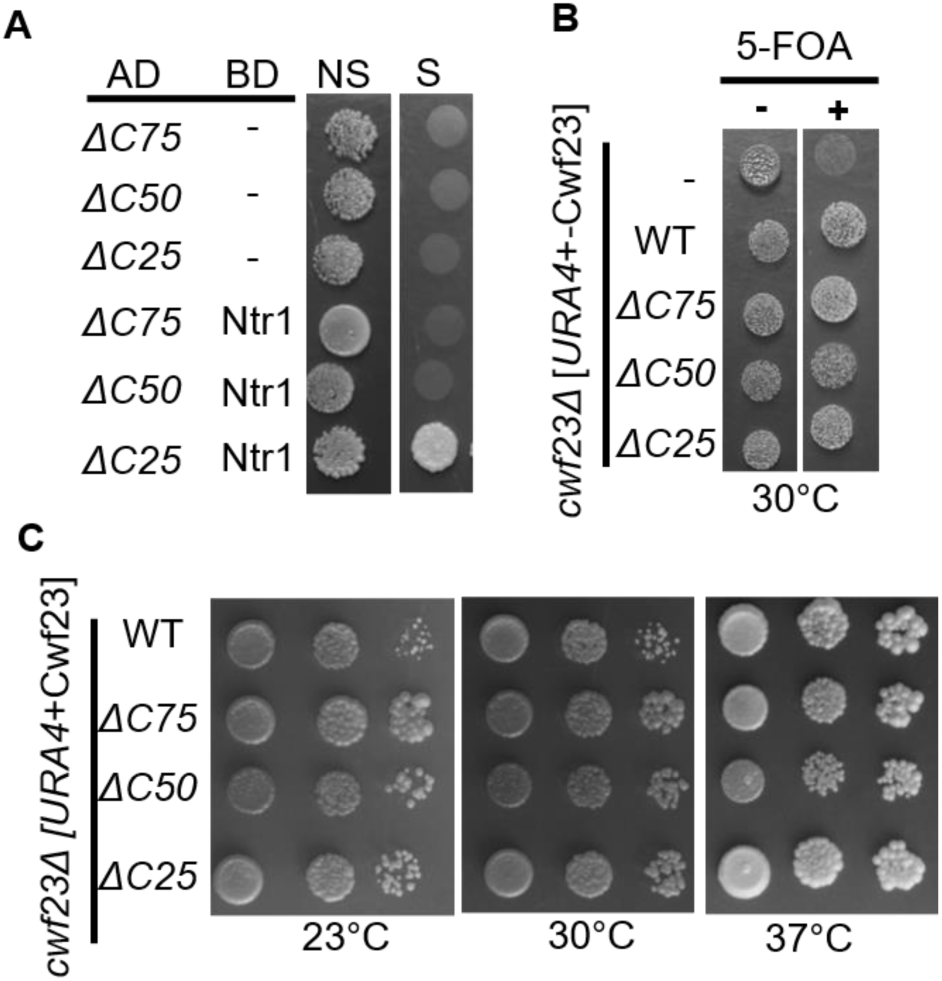
C-terminal region of Cwf23 is required for interaction with *Sp*Ntr1. **A.** Yeast two-hybrid interaction analysis of Cwf23 ΔC mutants with *Sp*Ntr1. Equal dilutions of two-hybrid tester strain (*AH109*) harboring *Sp*Ntr1 fused to the *GAL4*-binding domain (BD) and Cwf23 wildtype or the C-terminal truncation mutants fused to the *GAL4* activation domain (AD) or just the empty vectors (-) were spotted on non-selective (NS) or selective (S) medium. Plates were incubated at 30°C for 3 days. Growth on selective medium is indicative of a positive two-hybrid interaction. **B.** C-terminal of Cwf23 is not essential for cell viability. Yeast complementation analysis for Cwf23 *ΔC75*, *ΔC50* and *ΔC25*. Equal volumes of ten-fold serial dilutions of *cwf23Δ* cells harboring *URA3 CEN*-Cwf23 plasmid and pREP3x (*LEU*-based plasmid) empty vector (–), Cwf23 (WT) or ΔC mutants were spotted on media with or without 5-FOA and incubated at 30°C for 3 days. **C.** Growth assay of the Cwf23 mutants at different temperatures as indicated.

### J-domain of Cwf23 is essential

While the Cwc23 orthologs show significant divergence in their C-terminal region, the J-domain is conserved in all the identified orthologs (Raut et al., 2019). Interestingly, the J-domain of Cwc23 is completely dispensable for cell viability as well as pre-mRNA splicing in budding yeast (Sahi et al., 2010). In Cwf23, both the RRM and C-terminal *Sp*Ntr1-interating regions were dispensable. We hypothesized that in complex, “intron-rich” eukaryotes, the J-domain has more important roles to play than the simpler “intron-poor” eukaryotes, like budding yeast. To investigate the functional relevance of the Cwf23 J-domain *in vivo*, two independent mutants, *H37Q* and a complete deletion of the J-domain, *ΔJ* (1-66 amino acids), were generated. His37 is part of the invariable HPD motif in the J-domain of Cwf23 (Fig 4A). A Histidine-to-glutamic-acid substitution abolishes the ATPase stimulation ability of the JDPs and hence their Hsp70-dependent functions. Intriguingly, both the *H37Q* and *ΔJ* mutants were unable to complement the essential functions of Cwf23 *in vivo* (Fig 4B). To ensure that the non-complementation was not due to non-expression of mutant proteins, the mutants were sub-cloned with an N-terminal 1X HA-tag, followed by western analysis using anti-HA (12CA5, Sigma-Aldrich). Both the mutants were expressed and the levels were comparable with wild-type protein levels (Fig S1). To ascertain the Hsp70 co-chaperone activity of Cwf23, we tested the functionality of its J-domain by exploiting a previously established genetic approach. The temperature sensitivity of budding yeast cells lacking the multi-functional JDP, Ydj1, is significantly rescued by heterologous expression of J-domain fragments of many different JDPs (Sahi & Craig, 2007b; Tak et al., 2023; Verma et al., 2017). As expected, expression of wild-type Cwf23 J-domain fragment (1-165 amino acids) rescued the temperature sensitivity of *ydj1Δ* cells at 30°C while *H37Q* did not (Fig 4C). Further, using yeast two-hybrid, we analyzed if the J-domain of Cwf23 has any role in interaction with *Sp*Ntr1. Our results show that the J-domain is not required for Cwf23’s interaction with *Sp*Ntr1 (Fig 4D) as both *H37Q* and *ΔJ* mutants interacted with *Sp*Ntr1. Taken together, our results conclusively show that unlike Cwc23, the J-domain of Cwf23 is essential for its function *in-vivo* and the Hsp70-co-chaperone activity of Cwf23 is indispensable for cell viability.

**Figure 4.**
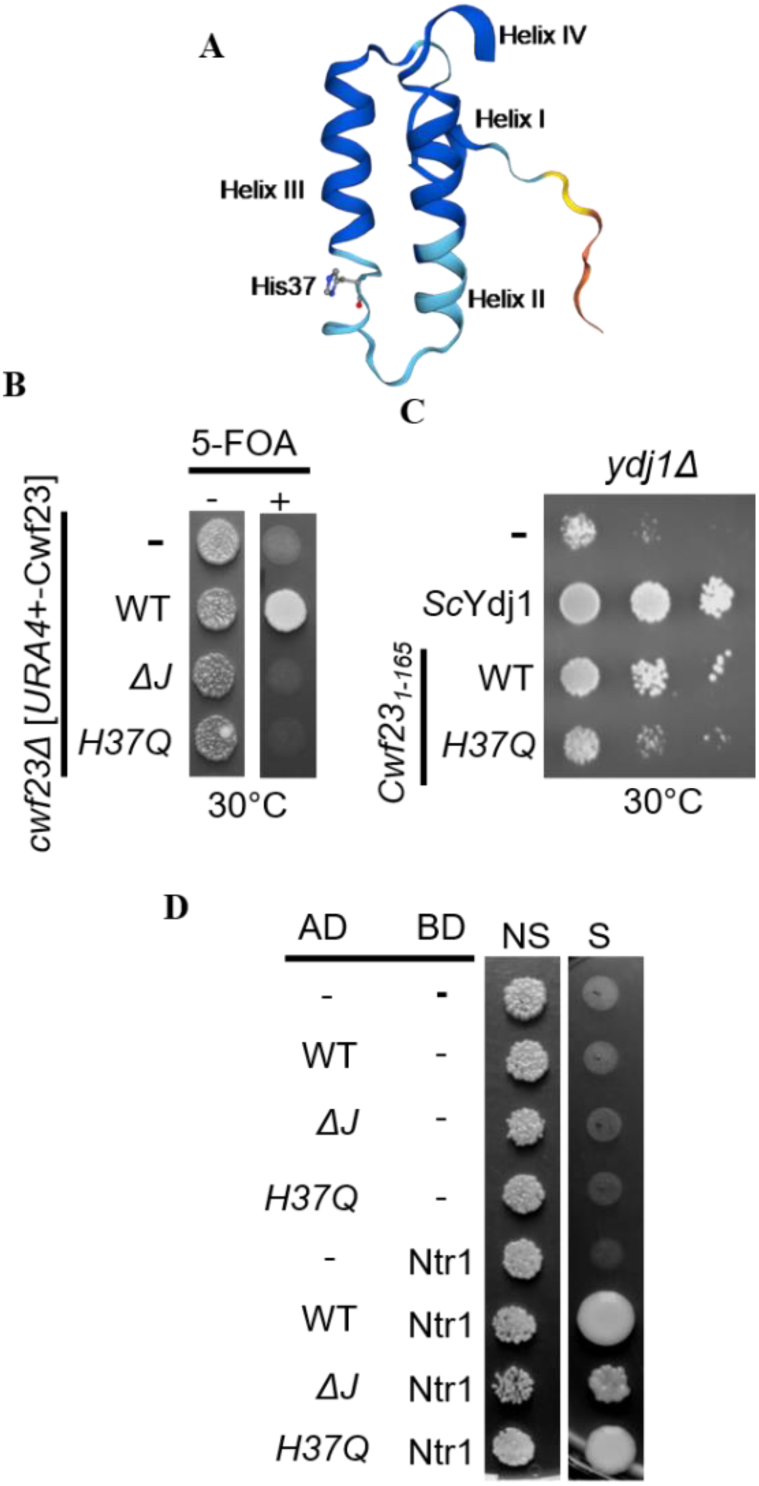
Cwf23 J-domain is indispensable for growth. **A.** Alpha fold structure of the J-domain showing alpha helices and histidine37 of Cwf23. **B.** Yeast complementation analysis for *Cwf23_ΔJ* and *H37Q* mutants. Equal volumes of ten-fold serial dilutions of *cwf23Δ* cells harboring *URA3 CEN*-Cwf23 plasmid and pREP3x (*LEU*-based plasmid) empty vector (–), Cwf23 (WT) or J domain mutants (*ΔJ* and *H37Q*) were spotted on media with or without 5-FOA and incubated at 30 °C for 3 days. **C.** *Cwf23_H37Q_* fails to rescue *ydj1Δ* cells. 10-fold serial dilutions of an equivalent number of *ydj1Δ* cells expressing *Sc*Ydj1, *Cwf23_1-165_* (WT), *Cwf23_1-165(H37Q)_* (*H37Q*) from the *TEF* promoter or harboring only an empty vector (-) were spotted on selective medium and incubated at 30°C for 3 days. **D.** Yeast two-hybrid interaction analysis of Cwf23 *ΔJ* and *H37Q* mutants with *Sp*Ntr1. Equal dilutions of two-hybrid tester strain (*AH109*) harboring *Sp*Ntr1 fused to the *GAL4*-binding domain (BD) and Cwf23 WT or the J-domain mutants (*ΔJ* and *H37Q*) fused to the *GAL4* activation domain (AD) or just the empty vectors (-) were spotted on non-selective (NS) or selective (S) medium. Plates were incubated at 30°C for 3 days. Growth on selective medium is indicative of a positive two-hybrid interaction.

### J-domain of Cwf23 is required for pre-mRNA splicing

Since both the J-domain mutants studied were non-viable, we set out to identify hypomorphic mutants in the Cwf23 J-domain. A multiple sequence alignment of Cwf23’s J-domain with other JDPs was performed to identify conserved, and potentially important amino acid residues within the J-domain of Cwf23, other than the invariant HPD (Fig S2). To more rigorously test the importance of Cwf23’s J-domain *in vivo* and pre-mRNA splicing, selected conserved amino acids in the Cwf23 J-domain (Fig 5A) were mutated to previously reported residues (Xue et al., 2018), and analyzed in *S. pombe* cells lacking Cwf23. Interestingly, all the mutants were lethal on SC and EMM plates, except *Y58N* (Fig 5B). Although viable, the *Y58N* mutant exhibited severe growth defects at 30°C as compared to the wild-type and could not form colonies at 37°C (Fig 5C).

**Figure 5.**
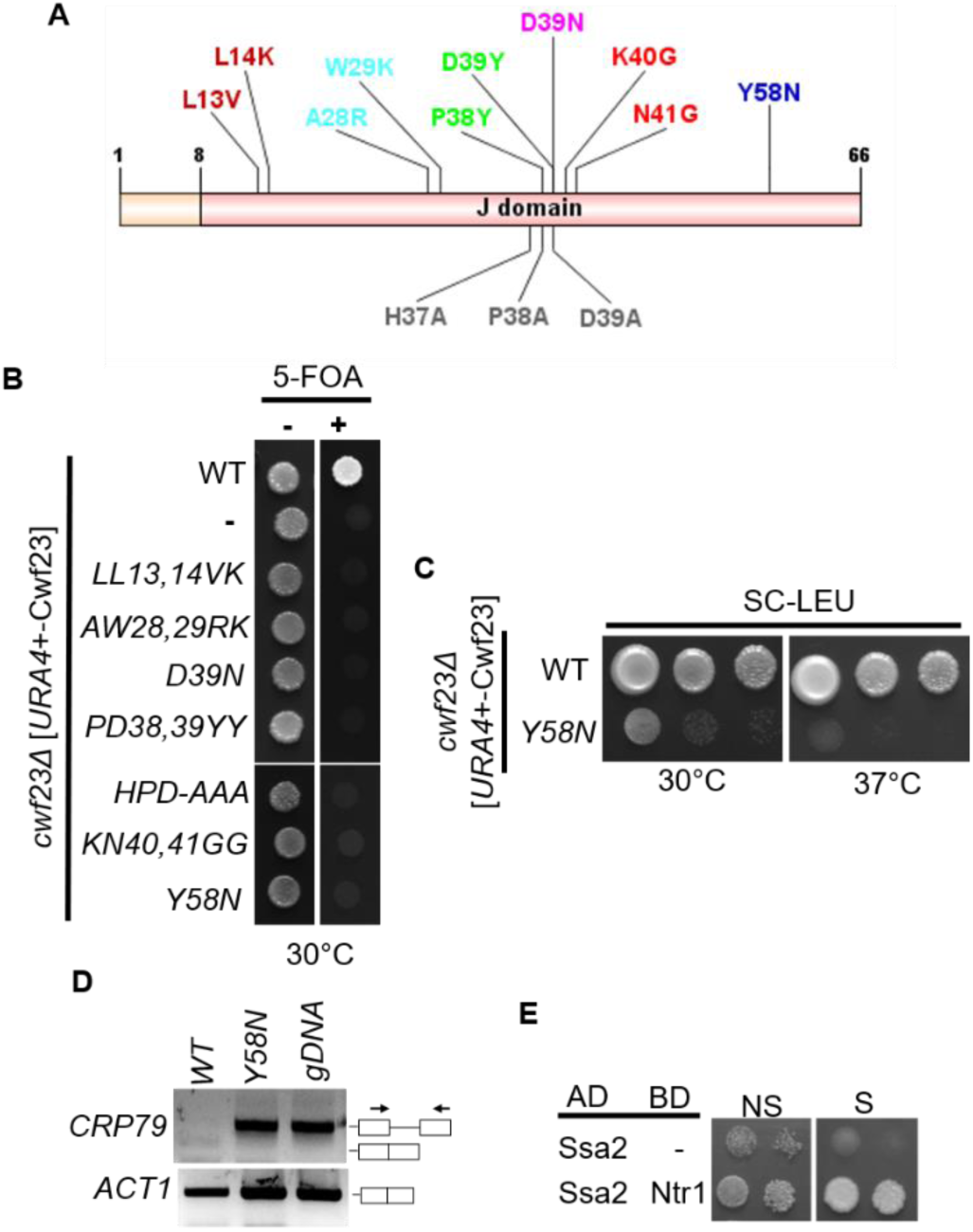
Cwf23 J-domain mutants, except *Y58N*, are lethal. **A.** Diagrammatic representation of the Cwf23 J domain mutants showing the positions and symbols of the changed residues (*LL13,14VK*; *AW28,29RK*; *D39N*; *HPD-AAA*; *PD28,39YY*; *KN40,41GG* and *Y58N*). **B.** J-domain mutants are inviable. Equal volume of ten-fold serial dilutions of *cwf23Δ* cells harboring *URA3 CEN*-Cwf23 plasmid and *LEU*-based plasmid with either no insert (–), endogenous wt Cwf23 (WT) or different mutants of Cwf23 were spotted on media with or without 5-FOA and incubated at 30 °C for 3 days. **C.** *Y58N* mutant is extremely sick but viable on extended incubation at 30°C. Equal volume of ten-fold serial dilutions of *cwf23Δ* cells harbouring wild-type Cwf23 (WT) or *Y58N* mutant of Cwf23 were spotted on SC-LEU media and incubated at the indicated temperatures (30°C and 37°C) for more than 7 days. **D.** Agarose gel pictures showing the expression and splicing pattern of CRP79. Total RNA from different strains was used to prepare cDNA. An equal quantity of cDNA was used for PCR with primers flanking the introns. Actin primers have been used to verify equal cDNA template for each strain (bottom panel). **E.** *Sp*Ntr1 interacts with *Sp*SSA2 in yeast two hybrid analysis. Equal dilutions of two-hybrid tester strain (*AH109*) harboring *Sp*Ntr1 or Ssa2 fused to the *GAL4*-binding domain (BD) and Cwf23, Prp43 and Ssa2 fused to the *GAL4* activation domain (AD) or just the empty vectors (-) were spotted on non-selective (NS) or selective (S) medium. Plates were incubated at 30°C for 3 days. Growth on selective medium is indicative of a positive two-hybrid interaction.

Finally, we asked whether the J-domain, and hence the Hsp70 co-chaperone activity of Cwf23 is also required for splicing. Out of the several genes that were analyzed for splicing defects above, a clear defect was noted in the CRP79 transcript in RRM mutants, so we checked the spliced and unspliced transcripts of CRP79 gene in the *Y58N* mutant background. Semi-quantitative RT-PCR data shows that CRP79 displayed a strong splicing defect in the Cwf23 *Y58N* mutant which was absent from the wild-type sample (Fig 5D). Our results show that J-domain is essential for Cwf23 function *in vivo* and defects in the J-domain affect pre-mRNA splicing in *S. pombe*.

### *Sp*Ntr1 interacts with *Sp*Ssa2

Although a direct involvement of an Hsp70 in spliceosomal remodelling is yet to be established, the data presented above does suggest that Cwf23 is cooperating with an Hsp70 in *S. pombe*. Although no Hsp70 has been reported to work with the NTR complex till date, the multi-functional Hsp70, Ssa2, was identified among proteins, pulled down by TAP-tagged Ntr1 (Cipakova et al., 2019). We tested if Ssa2 directly interacted with components of the NTR complex using yeast-two-hybrid. As shown in Fig 5E, Ssa2 interacted with *Sp*Ntr1 in our yeast two-hybrid analysis, evident from growth on selective media at 30°C. We could not detect a yeast-two-hybrid interaction of Ssa2 with either Cwf23 or Prp43 (*data not shown*). This suggests that Ssa2 could be the Hsp70, directly working with the NTR complex to mediate spliceosomal remodelling in fission yeast. Our results reinforce the idea of a direct involvement of the Hsp70-JDP machinery in the process of splicing where Ssa2 is the proposed Hsp70 in action.

## Discussion

Hsp70-JDP machinery is involved in various cellular functions including protein folding, refolding, transport across membranes, and formation and disassembly of protein complexes (Kampinga & Craig, 2010b; Mayer & Bukau, 2005; Qiu et al., 2006; Rosenzweig et al., 2019). Although highly likely, role of Hsp70-JDP machinery in spliceosome remodelling remains largely elusive. Cwc23, is a highly specialized JDP that works in spliceosome remodelling, through interaction with another essential spliceosomal remodelling factor, Ntr1 (Su et al., 2018; Tauchert et al., 2017; Tsai et al., 2005). Contrary to canonical JDPs, which require the J-domain to perform their functions, Cwc23 is unique, as its J-domain is dispensable for its essential function in budding yeast (Sahi et al., 2010). Interestingly, not only the J-domain in Cwc23 orthologs is conserved across species, Cwc23 orthologs in complex eukaryotes additionally have an RRM at their C-terminal. In the present study, we investigated the functional relevance of the RRM and J-domain of Cwf23, the Cwc23 ortholog in *S. pombe*. Our results show that although the RRM is important for Cwf23 functions *in vivo*, the requirement of the J-domain, hence the Hsp70 co-chaperone activity is consequential.

RRMs are one of the most abundant RNA binding domains (RBDs) known to engage in both protein-protein as well as RNA-protein interactions (Adam et al., 1986; Glisovic et al., 2008; Kielkopf et al., 2004; Lorković, 2009; Sachs et al., 1986). Cwc23 orthologs including *Hs*Cwc23, *Dm*Cwc23 and Cwf23, bind RNA *in vitro* (Raut et al., 2019). We found that RRM was required for optimal cell growth and pre-mRNA splicing. While the precise role of RRM in Cwc23 orthologs is yet to be established, a high throughput proteomic analysis of the human DnaJC17 interactome identified many splicing factors (Pascarella et al., 2018), suggesting a possibly broader role of RRM in establishing other, currently unknown protein/RNA interactions required for the functionality Cwc23 orthologs in complex eukaryotes. Nonetheless, while deletion of RRM affected RNA binding, *in vitro*, it had a relatively mild effect on growth. Moreover, *ΔRRM* exhibited defects in splicing of only a few transcripts analyzed. We propose that although not essential under physiological conditions, RRM may be required under stressful regimes. For example, in *S. cerevisiae*, the “otherwise” dispensable J-domain of Cwc23 becomes essential in the background of destabilizing mutations in disassembly factors, Ntr1 and Prp43 (Sahi et al., 2010). While we cannot rule out the possibility that RRM is required for interaction with RNA, protein or both on the spliceosome, our results endorse an assumption that Cwf23’s RRM is rather accessory and not essential in *S. pombe* under physiological conditions.

The C-terminal of Cwf23 was required for interaction with *Sp*Ntr1 as *ΔC50* and *ΔC75* truncations affected interaction with *Sp*Ntr1 in yeast two-hybrid assays. However, none of these mutants affected cell growth at any of the temperatures tested. This is in contrast to budding yeast, where defects in Cwc23:Ntr1 interaction result in severe growth and global pre-mRNA splicing defects (Sahi et al., 2010). Additionally, Cwc23 is required for maintaining Ntr1 levels on the spliceosome. Depletion of Cwc23 also depleted Ntr1 from splicing extracts of *S. cerevisiae* and hampered spliceosome disassembly (Su et al., 2018), indicating a role of Cwc23 in stabilizing Ntr1. While the intra-species interaction between Cwf23 and Ntr1 orthologs is conserved (Raut et al., 2019), and Cwf23 C-term is the primary site of contact between Cwf23 and *Sp*Ntr1, possibly additional contacts, may be stabilizing this interaction *in vivo*. Indeed, identification of Ssa2 as an interactor of *Sp*Ntr1 in our yeast two-hybrid assays supports the hypothesis that Cwf23 may be involved in a bipartite interaction with *Sp*Ntr1, one directly through its C-term and another via the Hsp70, Ssa2.

Dispensability of Cwf23 C-term and RRM and identification of the Hsp70, Ssa2 as a direct interactor of *Sp*Ntr1 prompted us to investigate the importance of its J-domain, which interact with Hsp70s and define the functionality of a JDP. A J-domain fragment, lacking most of the C-term, was functional as it could efficiently rescue the temperature sensitivity of *ydj1Δ* strain of budding yeast. However, unlike Cwc23, the J-domain of Cwf23 is essential for cell viability suggesting that the Hsp70-dependent function of Cwf23 defines the functionality of this JDP in *S. pombe*. Both the deletion and changes in the HPD motif rendered the J-domain completely inactive. Interestingly, several other point mutations (other than the critical HPD) within the J-domain of Cwf23 also resulted in lethality. The same mutations in the J-domain of other JDPs do not affect J-domain function drastically (Jiang et al., 2007). One of these mutants, *Y58N*, rescued the essentiality of *cwf23Δ,* however was extremely sick and exhibited RNA splicing defect. Our results suggest that the Hsp70-co-chaperone functions of Cwf23 are critical and perturbations in the functionality of the J-domain lead to severe effects on cell viability. In budding yeast, Hsp70 and Hsp104 restore splicing in wild-type strains following heat stress by reactivation of heat-denatured spliceosomal proteins (Vogel et al., 1995; Yost & Lindquist, 1986). The idea of an Hsp70 dependent role of DnaJC17 in splicing has been recently proposed in human cells (Allegakoen et al., 2023, bioRxiv preprint) but the evidence for a direct involvement of an Hsp70 with splicing machinery is still lacking. The fact that the J-domain of Cwf23 plays an essential role, hints towards a direct involvement of an Hsp70-JDP machinery in the spliceosome remodeling in intron-rich eukaryotes.

In summary, we assert that while the proper functioning of Cwf23 requires its RRM, the J-domain is indispensable *in vivo* (Fig 6). The crucial role played by the J-domain underscores the involvement of the Hsp70-JDP system in the remodelling of the spliceosome, especially in intron-rich eukaryotes. Although speculative at the moment, our results posit that a bipartite interaction between Cwf23 and *Sp*Ntr1 may be important for the stability and functionality of the NTR complex in fission yeast (Fig 6). Whether a similar system prevails in other intron-rich eukaryotes, or additional layers of complexity exist, needs to be addressed. Investigating the nuanced functions of Cwf23 and other Cwc23 orthologs in plants and metazoans becomes particularly valuable, given the complexity of the splicing system in these organisms, demanding precision and efficiency in the vital process of splicing. While the Hsp70-JDP machinery is acknowledged for its role in remodeling various protein complexes, our findings bring us closest to identifying the co-partners of this chaperone system in the process of spliceosomal disassembly.

**Figure 6.**
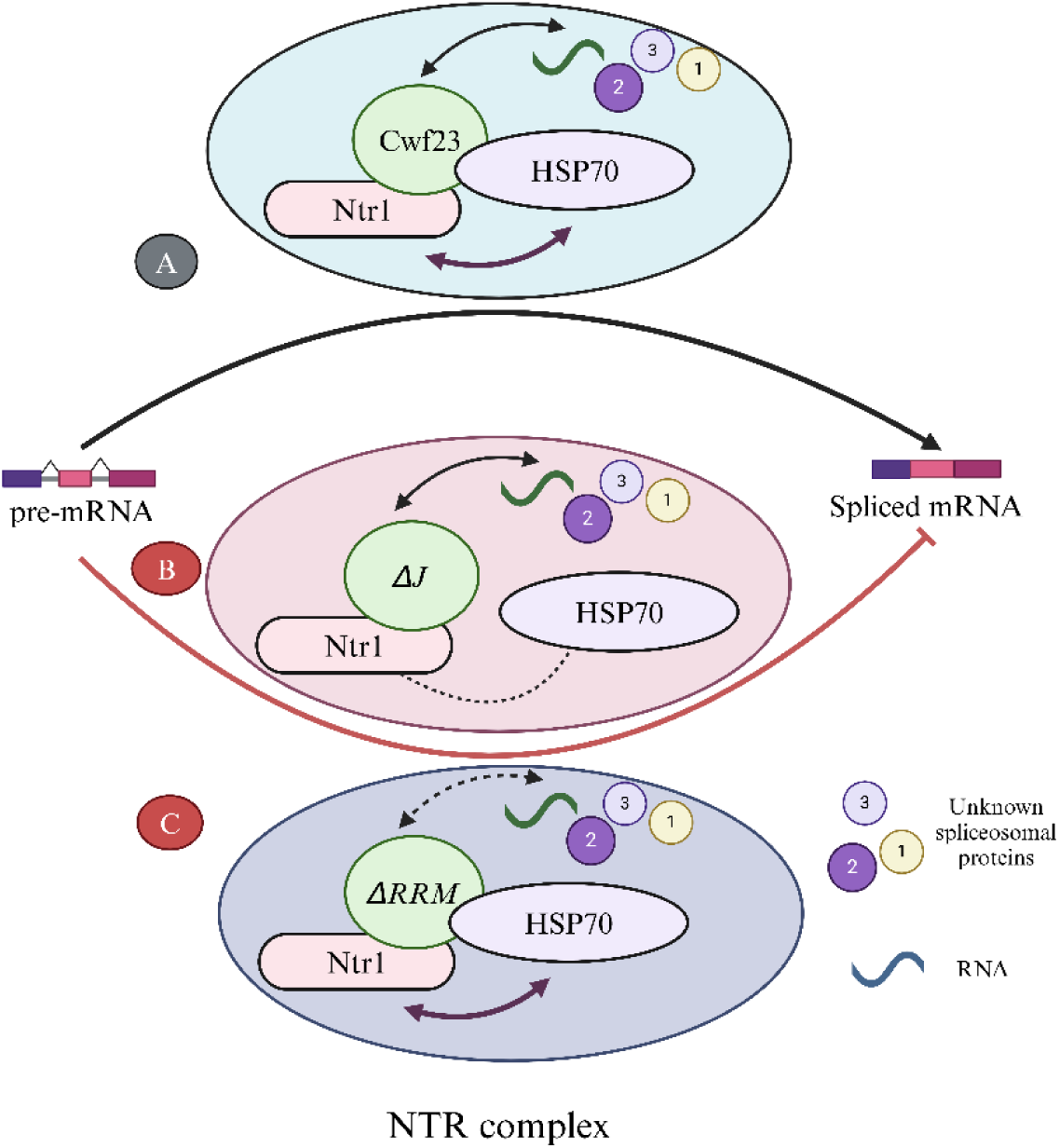
Working model for the role of Cwf23 in spliceosome disassembly in *S. pombe*. **A.** Cwf23 interacts with Ntr1 and possibly with unknown splicing factors/ RNA through its RRM. Ntr1 also interacts with Hsp70 Ssa2, which is required for proper functioning of the NTR complex. **B.** *ΔJ*, which is defective in interaction with Hsp70, may also affect the Ntr1:Hsp70 interaction *in vivo*. Additionally, *ΔJ* may directly affect a yet unknown chaperone function of Hsp70 on the spliceosome, thereby impacting cell growth and pre-mRNA splicing. **C.** *ΔRRM* affects Cwf23’s interaction with some unknown splicing factors and/or RNAs, however it only mildly affects growth and splicing as interactions between Ntr1, Cwc23 and Hsp70 are not compromised.

## Acknowledgements

This work was supported by funding from the Department of Biotechnology (BT/PR12149/BRB/10/1348/2014) and intramural funds from IISER Bhopal to CS. KY, SR and MGR thank CSIR, GATE and UGC respectively for the fellowship. We thank CS Lab members for their discussion and insightful suggestions.

## Accession Numbers

The sequence information of the genes examined in this study was sourced from the PomBase. The amino acid sequences were sourced from SGD, PomBase, TAIR and UniProt. The accession numbers for these genes are as follows:

Cwf23 (SPCC10H11.02), *Sp*Ntr1 (SPAC1486.03c), *Sp*Ssa2 (SPCC1739.13), *Sc*Cwc23 (YGL128C), *Sc*Ydj1(YNL064C), *Sc*Caj1 (YER048C), *Sc*Jjj1 (YNL227C), *Sc*Sis1 (YNL007C), *At*Cwc23(AT5G23590), *Hs*Cwc23 (DNAJC17), *Dm*Cwc23 (CG17187), *Ec*DnaJ (P08622)

## Author contributions

CS and SR conceived and initiated the project; CS and SKM supervised the research and furnished laboratory facilities and funding; KY and CS designed the experiments and KY performed most of the experiments and analyzed the data; MGR, GS, MD and RB contributed to the generation of valuable resources, data acquisition and analysis. KY, MGR and CS wrote the manuscript.

## Supplemental data

Supplemental Table S2. List of genes used for checking splicing defects in this study.

Supplemental Table S1. List of constructs used in this study.

Supplemental Figure S1: Western blot analysis of different Cwf23 mutants analyzed in this study.

Supplemental Figure S2. Sequence alignment of J-domain fragments of Cwc23 orthologs.

## Data availability

The authors confirm that all experimental data are available and accessible via the main text and/or the supplemental data.

