## Supplemental figures and tables for "Inclusion of Hsp70 co-chaperone and RNA binding activities facilitate optimal function of spliceosomal disassembly factor Cwf23 in intron-rich *Schizosaccharomyces pombe*"

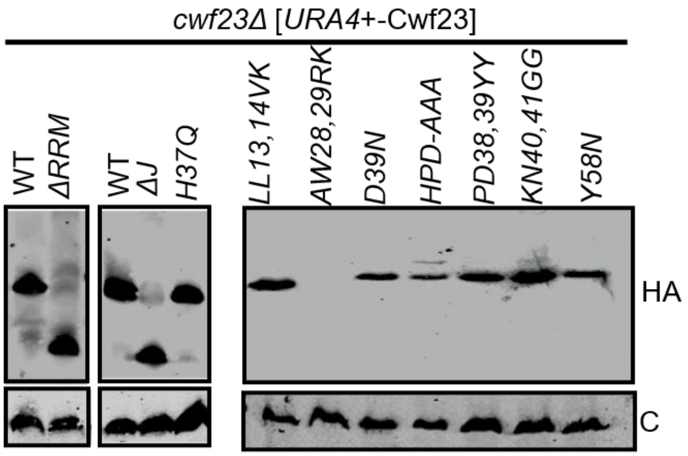


**Figure S1. Western blot analysis to show comparable expression of Cwf23 WT and mutants.** Equal amounts of total cell lysate prepared from WT and *cwf23Δ* strains transformed with plasmids carrying different versions of Cwf23 (WT and different mutants) were resolved on 12.5% SDS-PAGE, electroblotted onto a PVDF membrane and probed with anti-HA and anti- rabbit secondary antibody. The blot was visualized by chemiluminescence on LI-COR Odyssey 9120. Anti-Tubulin was used as a loading control.

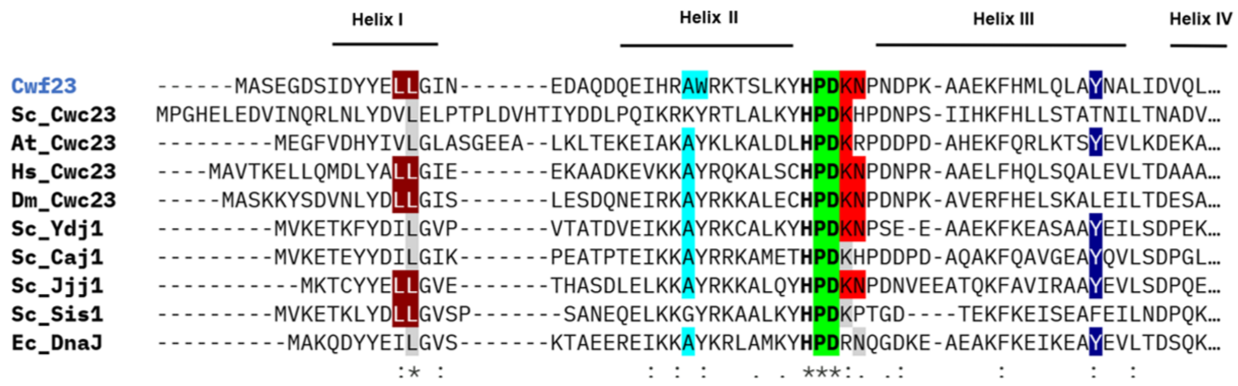


**Figure S2. Sequence alignment of J-domain fragments of Cwc23 orthologs.** Multiple sequence alignment of Cwf23 orthologs: *Sc*Cwc23 (1-80), *At*Cwc23 (1-68), *Hs*Cwc23 (1-68), *Dm*Cwc23 (1-67) and other Yeast JDPs: *Sc*Ydj1 (1-62), *Sc*Caj1 (1-63) , *Sc*Jjj1 (1-62) , *Sc*Sis1 (1-60), *Ec*DnaJ (1-62) was carried out using Clustal Omega (https://www.ebi.ac.uk/jdispatcher/msa/clustalo) (Madeira F et al,. 2022). The four alpha helices of the J-domain are shown above their corresponding sequences.

**Supplemental Table S1. List of genes used for checking splicing defects in this study.**

| Gene | Description |
|---|---|
| MMS1 | Cul8-RING ubiquitin ligase complex WD repeat protein subunit Mms1 |
| PST2 | Clr6 histone deacetylase complex subunit Pst2 |
| RHP23 (Rad23 homolog Rhp23) | Proteasome-associated ubiquitin receptor |
| CDK9 | Cyclin-dependent protein kinase |
| CRP79 | Poly(A) binding protein Crp79 |

**Supplemental Table S2. List of plasmids generated and/or used in this study.**

| Plasmid ID | Gene | Base Vector | Source |
|---|---|---|---|
| pCS889 | *Cwf23* | pREP3X | This study |
| pCS1231 | *Cwf23_ΔRRM* | pREP3X | This study |
| pCS1402 | HA _*Cwf23* | pREP3X | This study |
| pCS1404 | HA_*Cwf23_ΔRRM* | pREP3X | This study |
| pCS1767 | *Cwf23_ΔC75* | pREP3X | This study |
| pCS1768 | *Cwf23_ΔC50* | pREP3X | This study |
| pCS1921 | *Cwf23_ΔC25* | pREP3X | This study |
| pCS1771 | HA_*Cwf23_ΔC75* | pREP3X | This study |
| pCS1770 | HA_*Cwf23_ΔC50* | pREP3X | This study |
| pCS1769 | HA_*Cwf23_ΔC25* | pREP3X | This study |
| pCS242 | *Cwf23_ ΔJ* | pREP3X | This study |
| pCS1151 | *Cwf23_ H37Q* | pREP3X | This study |
| pCS1403 | HA_*Cwf23_ ΔJ* | pREP3X | This study |
| pCS1922 | HA_*Cwf23_ H37Q* | pREP3X | This study |
| pCS1912 | HA_*Cwf23_LL13,14VK* | pREP3X | This study |
| pCS1913 | HA_*Cwf23_AW28,29RK* | pREP3X | This study |
| pCS1914 | HA_*Cwf23_D39N* | pREP3X | This study |
| pCS1915 | HA_*Cwf23_HPD-AAA* | pREP3X | This study |
| pCS1916 | HA_*Cwf23_PD28,39YY* | pREP3X | This study |
| pCS1917 | HA_*Cwf23_KN40,41GG* | pREP3X | This study |
| pCS1918 | HA_*Cwf23_Y58N* | pREP3X | This study |
| pCS661 | *Cwf23* | pDEST-pGADT7 | Raut et al. (2018) |
| pCS1237 | *Cwf23_ΔRRM* | pDEST-pGADT7 | This study |
| pCS1772 | *Cwf23_ΔC75* | pDEST-pGADT7 | This study |

| | | | |
|---|---|---|---|
| pCS1773 | *Cwf23_ΔC50* | pDEST-pGADT7 | This study |
| pCS1919 | *Cwf23_ΔC25* | pDEST-pGADT7 | This study |
| pCS1152 | *Cwf23_ ΔJ* | pDEST-pGADT7 | This study |
| pCS1234 | *Cwf23_ H37Q* | pDEST-pGADT7 | This study |
| pCS1789 | *Sp_Ssa2* | pDEST-pGADT7 | This study |
| pCS845 | *Sp_Ntr1* | pDEST-pGBKT7 | Raut et al. (2018) |
| pCS404 | *Cwf23* | pGEX6P1 | Raut et al. (2018) |
| pCS1920 | *Cwf23_ ΔRRM* | pGEX6P1 | This study |
| pCS1821 | *Cwf23_1-165* | pRS414 *TEF* | This study |
| pCS1823 | *Cwf23_1-165_H37Q* | pRS414 *TEF* | This study |
| pCS167 | *Sc_Ydj1* | pRS414 *TEF* | Sahi, C. and Craig, E.A. (2007) |
